## Supplementary material for "Dual Recognition of Multiple Signals in Bacterial Outer Membrane Proteins Enhance Assembly and Maintain Membrane Integrity": figure supplements

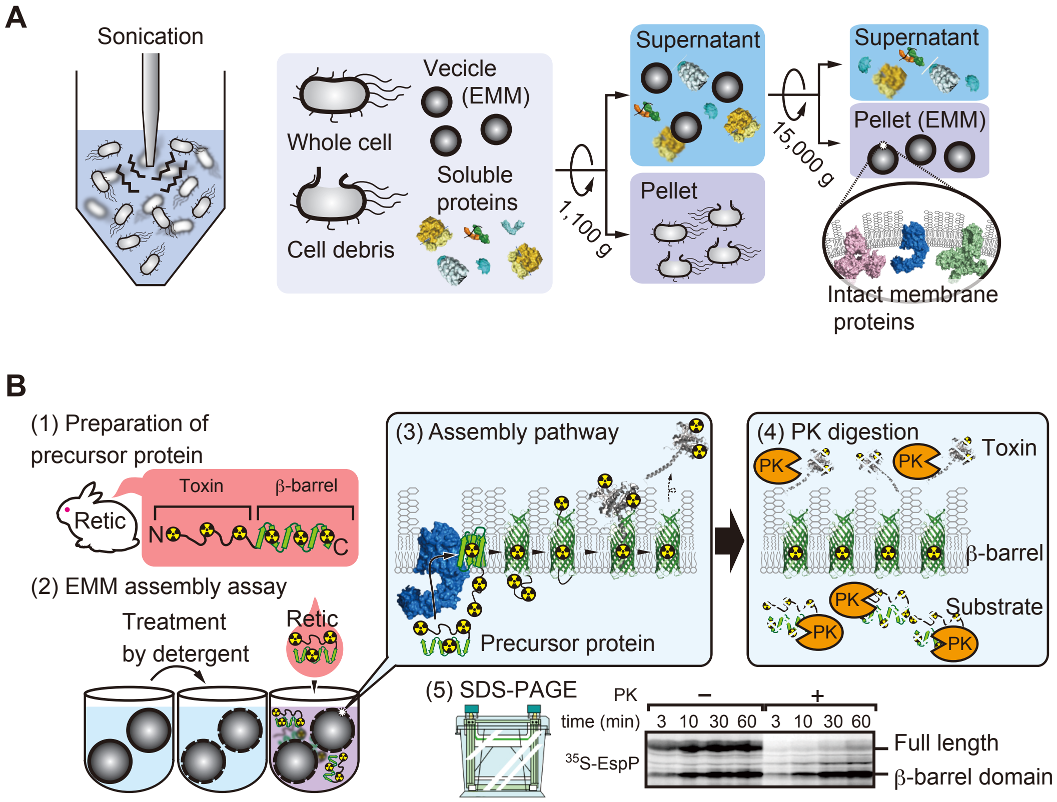
A
Sonication
Whole cell
Vecicle
(EMM)
Soluble
proteins
Cell debris
1,100 g
Supernatant
Pellet
15,000 g
Supernatant
Pellet (EMM)
Intact membrane
proteins
B
(1) Preparation of
precursor protein
Retic
Toxin
β-barrel
N
C
(2) EMM assembly assay
Treatment
by detergent
Retic
(3) Assembly pathway
Precursor protein
(4) PK digestion
PK
Toxin
β-barrel
Substrate
(5) SDS-PAGE
PK
–
+
time (min)
3 10 30 60 3 10 30 60
35S-EspP
Full length
β-barrel domain

### *E. coli* OmpC sequence

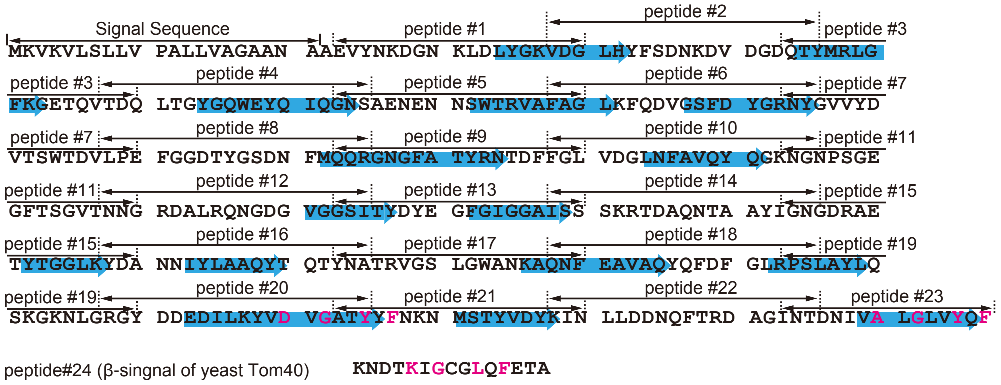

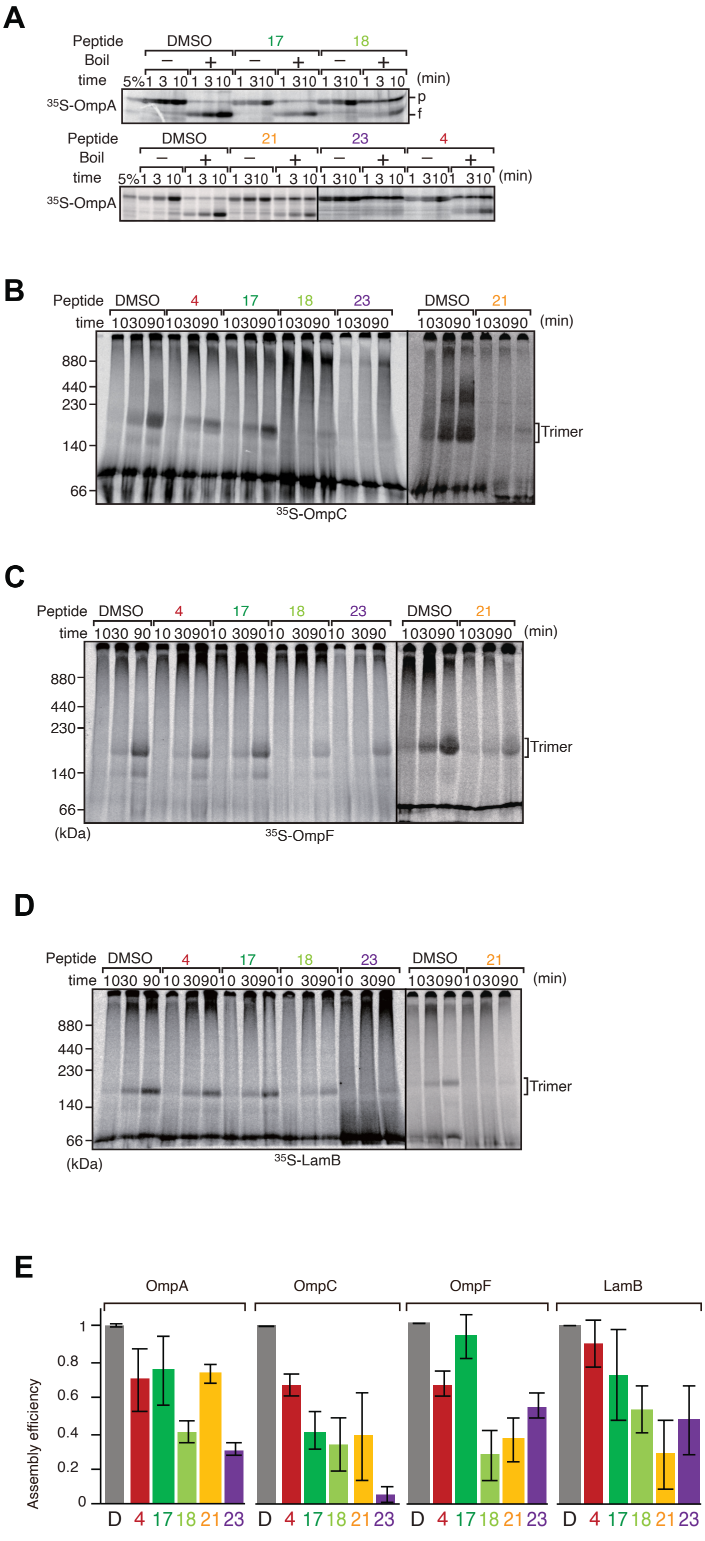
A
Peptide DMSO 17 18
Boil − + − + − +
time 5% 1 3 10 1 3 10 1 3 10 1 3 10 1 3 10 1 3 10 (min)
35S-OmpA
p
f
Peptide DMSO 21 23 4
Boil − + − + − + − +
time 5% 1 3 10 1 3 10 1 3 10 1 3 10 1 3 10 1 3 10 1 3 10 1 3 10 (min)
35S-OmpA
B
Peptide DMSO 4 17 18 23 DMSO 21
time 10 30 90 10 30 90 10 30 90 10 30 90 10 30 90 10 30 90 10 30 90 (min)
880
440
230
140
66
Trimer
35S-OmpC
C
Peptide DMSO 4 17 18 23 DMSO 21
time 10 30 90 10 30 90 10 30 90 10 30 90 10 30 90 10 30 90 10 30 90 (min)
880
440
230
140
66
(kDa)
Trimer
35S-OmpF
D
Peptide DMSO 4 17 18 23 DMSO 21
time 10 30 90 10 30 90 10 30 90 10 30 90 10 30 90 10 30 90 10 30 90 (min)
880
440
230
140
66
(kDa)
Trimer
35S-LamB
E
OmpA OmpC OmpF LamB
Assembly efficiency
1
0.8
0.6
0.4
0.2
0
D 4 17 18 21 23 D 4 17 18 21 23 D 4 17 18 21 23 D 4 17 18 21 23

A

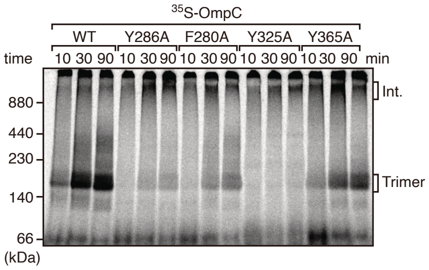

35S-OmpC
WT
Y286A
F280A
Y325A
Y365A
time
10 30 90 10 30 90 10 30 90 10 30 90 10 30 90 min
Int.
Trimer
880
440
230
140
66
(kDa)

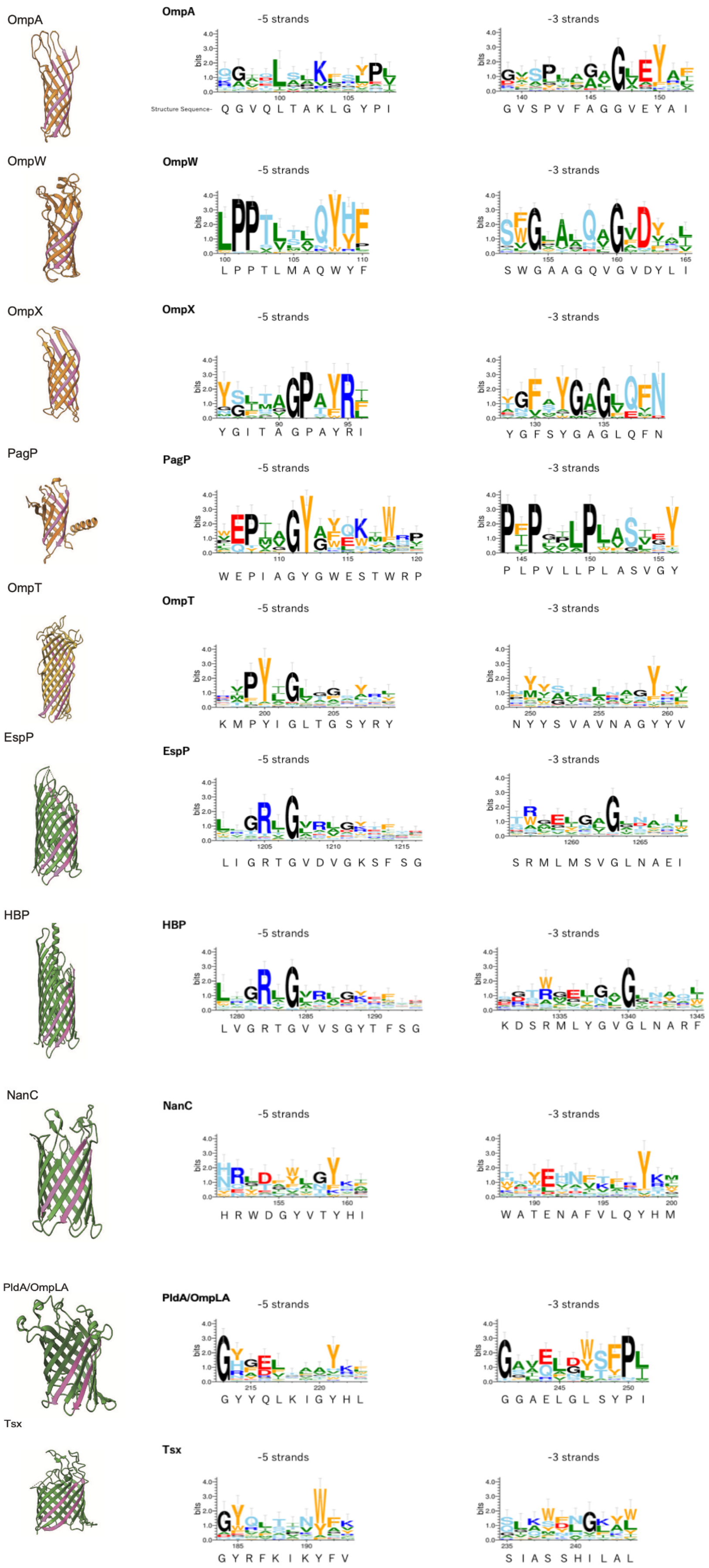

OmpA
-5 strands
-3 strands
Structure Sequence- Q G V Q L T A K L G Y P I
G V S P V F A G G V E Y A I
OmpW
-5 strands
-3 strands
L P P T L M A Q W Y F
S W G A A G Q V G V D Y L I
OmpX
-5 strands
-3 strands
Y G I T A G P A Y R I
Y G F S Y G A G L Q F N
PagP
-5 strands
-3 strands
W E P I A G Y G W E S T W R P
P L P V L L P L A S V G Y
OmpT
-5 strands
-3 strands
K M P Y I G L T G S Y R Y
N Y Y S V A V N A G Y Y V
EspP
-5 strands
-3 strands
L I G R T G V D V G K S F S G
S R M L M S V G L N A E I
HBP
-5 strands
-3 strands
L V G R T G V V S G Y T F S G
K D S R M L Y G V G L N A R F
NanC
-5 strands
-3 strands
H R W D G Y V T Y H I
W A T E N A F V L Q Y H M
PldA/OmpLA
-5 strands
-3 strands
G Y Y Q L K I G Y H L
G G A E L G L S Y P I
Tsx
-5 strands
-3 strands
G Y R F K I K Y F V
S I A S S H I L A L

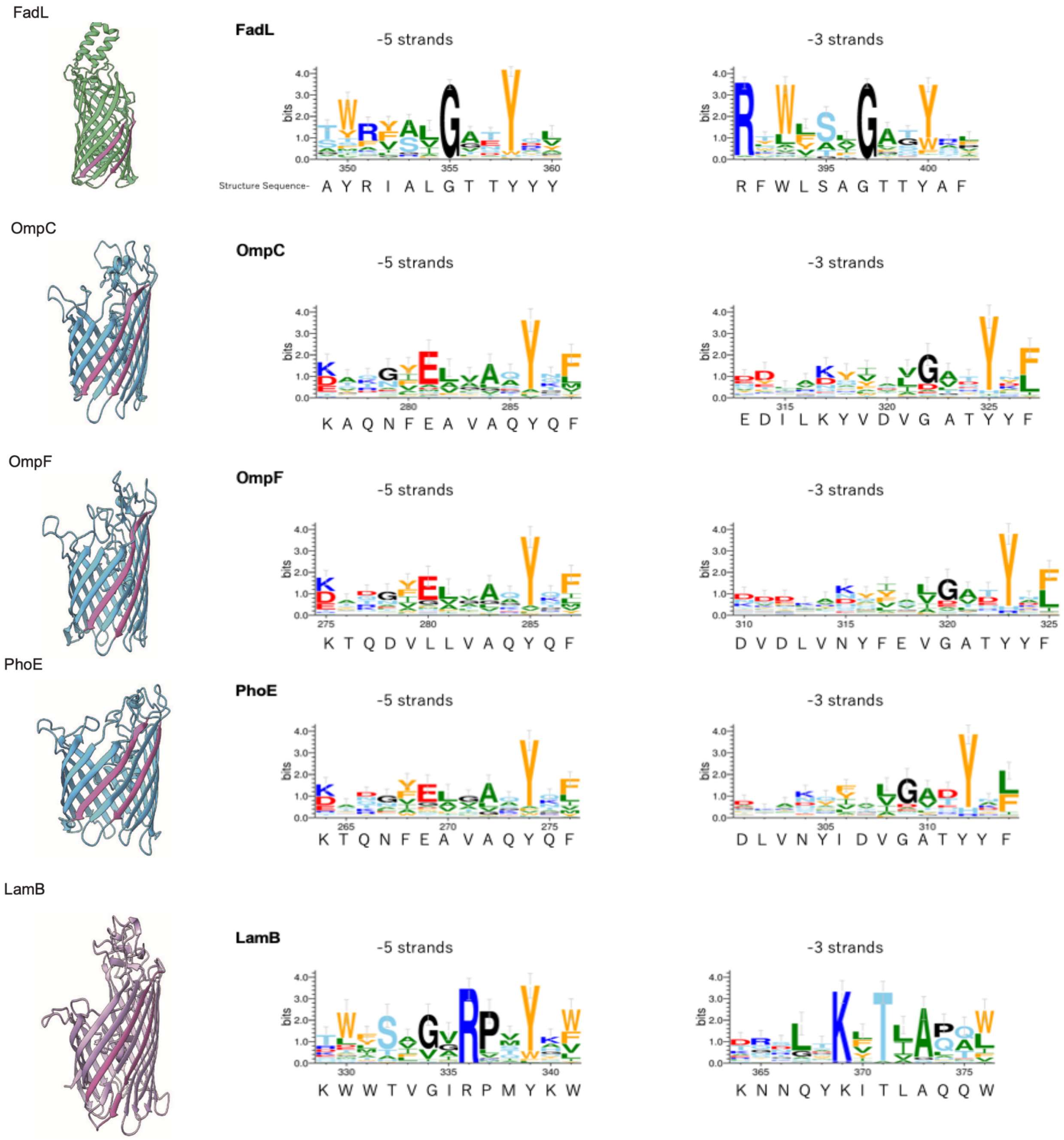
FadL
FadL
-5 strands
-3 strands
bits
Structure Sequence- A Y R I A L G T T Y Y Y
R F W L S A G T T Y A F
OmpC
OmpC
-5 strands
-3 strands
K A Q N F E A V A Q Y Q F
E D I L K Y V D V G A T Y Y F
OmpF
OmpF
-5 strands
-3 strands
K T Q D V L L V A Q Y Q F
D V D L V N Y F E V G A T Y Y F
PhoE
PhoE
-5 strands
-3 strands
K T Q N F E A V A Q Y Q F
D L V N Y I D V G A T Y Y F
LamB
LamB
-5 strands
-3 strands
K W W T V G I R P M Y K W
K N N Q Y K I T L A Q Q W

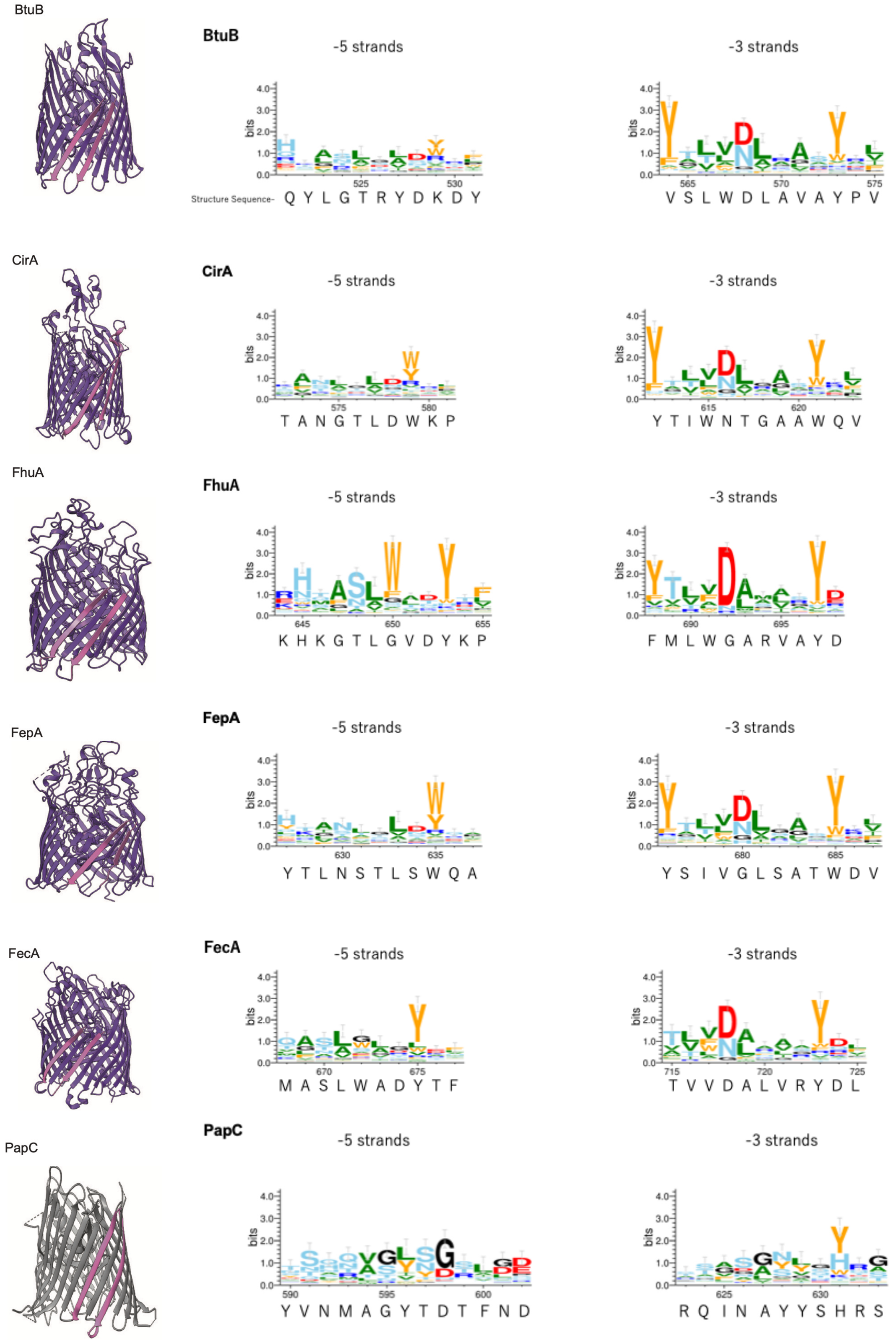
BtuB
-5 strands
-3 strands
Structure Sequence-
Q Y L G T R Y D K D Y
V S L W D L A V A Y P V
CirA
-5 strands
-3 strands
T A N G T L D W K P
Y T I W N T G A A W Q V
FhuA
-5 strands
-3 strands
K H K G T L G V D Y K P
F M L W G A R V A Y D
FepA
-5 strands
-3 strands
Y T L N S T L S W Q A
Y S I V G L S A T W D V
FecA
-5 strands
-3 strands
M A S L W A D Y T F
T V V D A L V R Y D L
PapC
-5 strands
-3 strands
Y V N M A G Y T D T F N D
R Q I N A Y Y S H R S

**A**

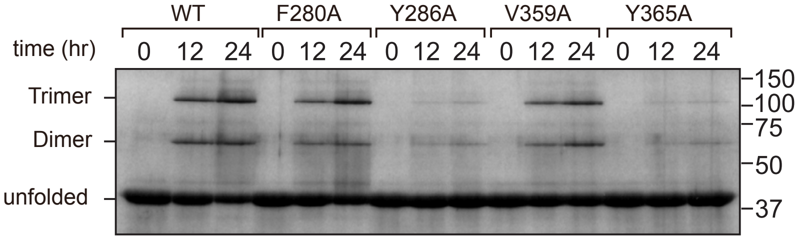


Non-Boiled (25°C)

**B**

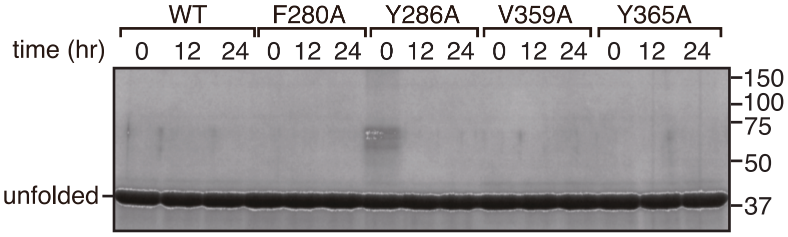


Boiled (95°C)

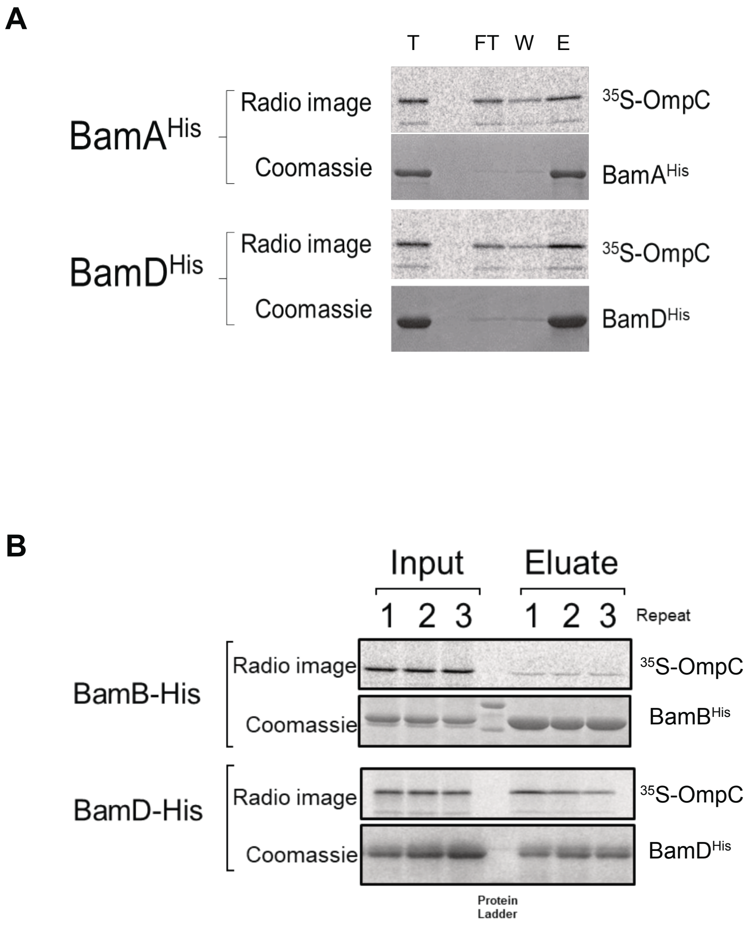
A
T
FT
W
E
BamA^His
Radio image
Coomassie
35S-OmpC
BamA^His
BamD^His
Radio image
Coomassie
35S-OmpC
BamD^His
B
Input
Eluate
1 2 3
1 2 3
Repeat
BamB-His
Radio image
Coomassie
35S-OmpC
BamB^His
BamD-His
Radio image
Coomassie
35S-OmpC
BamD^His
Protein Ladder

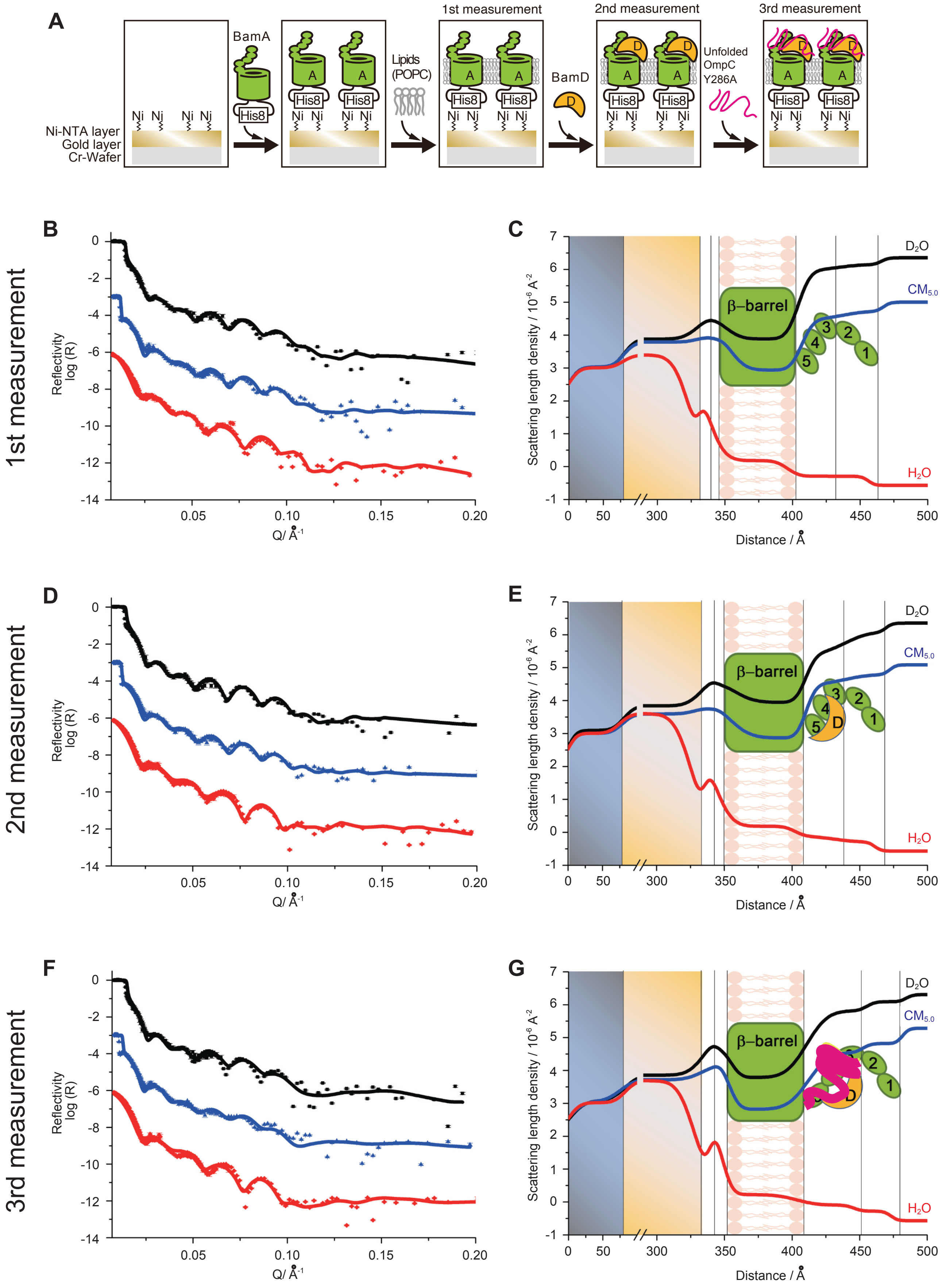
A
Ni-NTA layer
Gold layer
Cr-Wafer
Ni Ni Ni Ni
BamA
His8
A
Lipids (POPC)
1st measurement
BamD
D
2nd measurement
Unfolded OmpC Y286A
3rd measurement
B
1st measurement
Reflectivity log (R)
Q/ Å-1
C
Scattering length density / 10-6 Å-2
Distance / Å
β–barrel
D2O
CM5.0
H2O
D
2nd measurement
E
F
3rd measurement
G

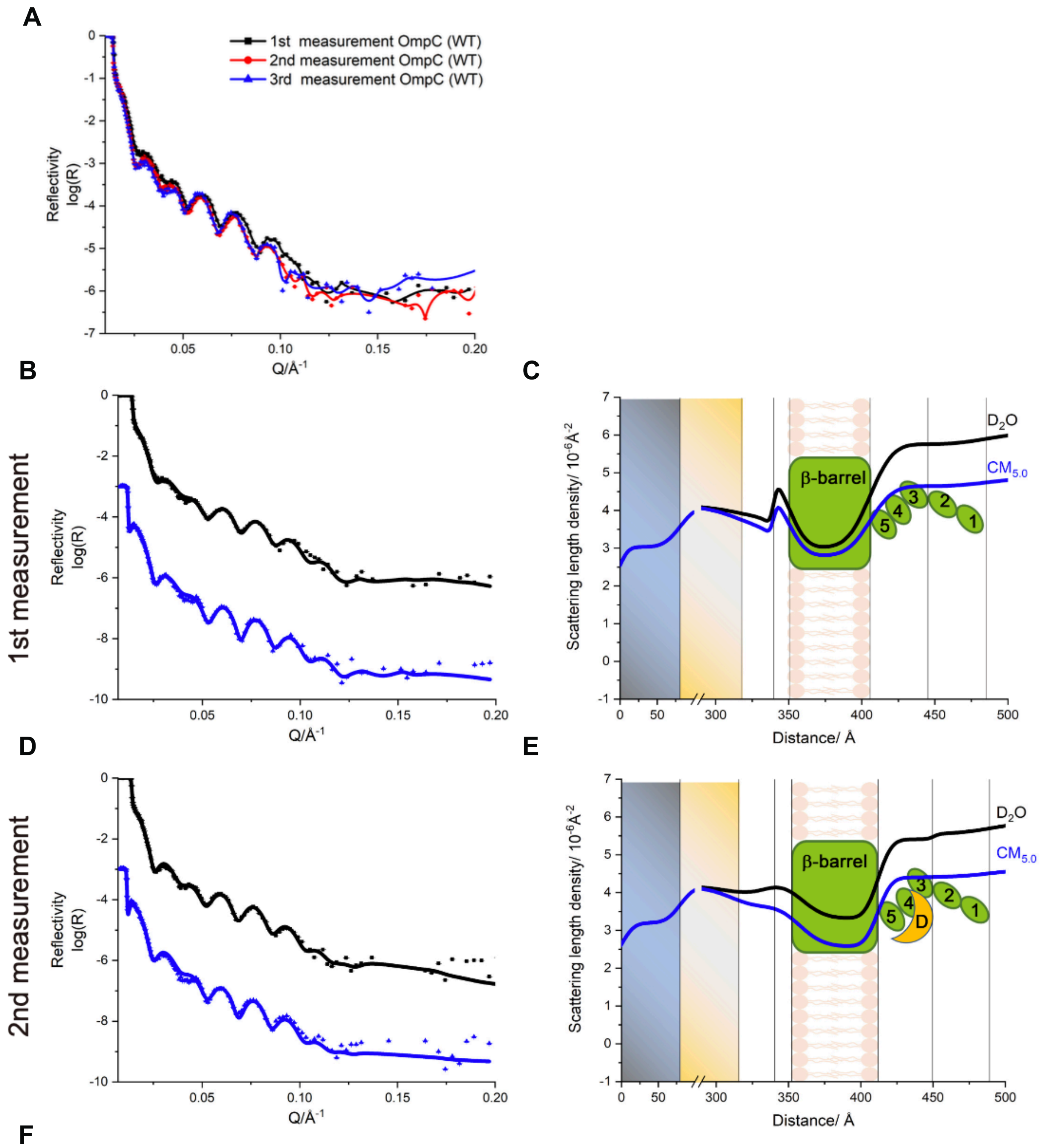


F

1st measurement 2nd measurement 3rd measurement (no change compared to 2nd measurement)

| Layers | Thickness (Å) | Volume fraction (%) | | | | Roughness (Å) |
|---|---|---|---|---|---|---|
| | | BamA | POPC | BamD | Solution | |
| His8 | 10.8±0.7 11.8±1.0 | - - | - - | - | 32.8±4.5 33.2±0.7 | 2.1±1.1 13.4±0.4 |
| β-Barrel | 55.2±0.2 59.6±0.4 | 33.0±1.5 33.0±1.8 | 35.6±2.9 34.1±0.6 | - | 31.4±2.9 45.0±1.9 | 9.2±0.8 15.1±0.5 |
| P3-5 | 39.7±0.4 37.8±0.8 | 11.3±0.1 11.3±0.1 | - - | 17.8±1.0 | 88.7±0.4 70.9±1.0 | 12.3±0.2 7.5±0.3 |
| P1-2 | 39.7±0.4 39.4±0.7 | 11.3±0.1 11.3±0.2 | - - | - | 88.7±0.5 88.7±0.2 | 12.0±6.3 7.5±3.5 |

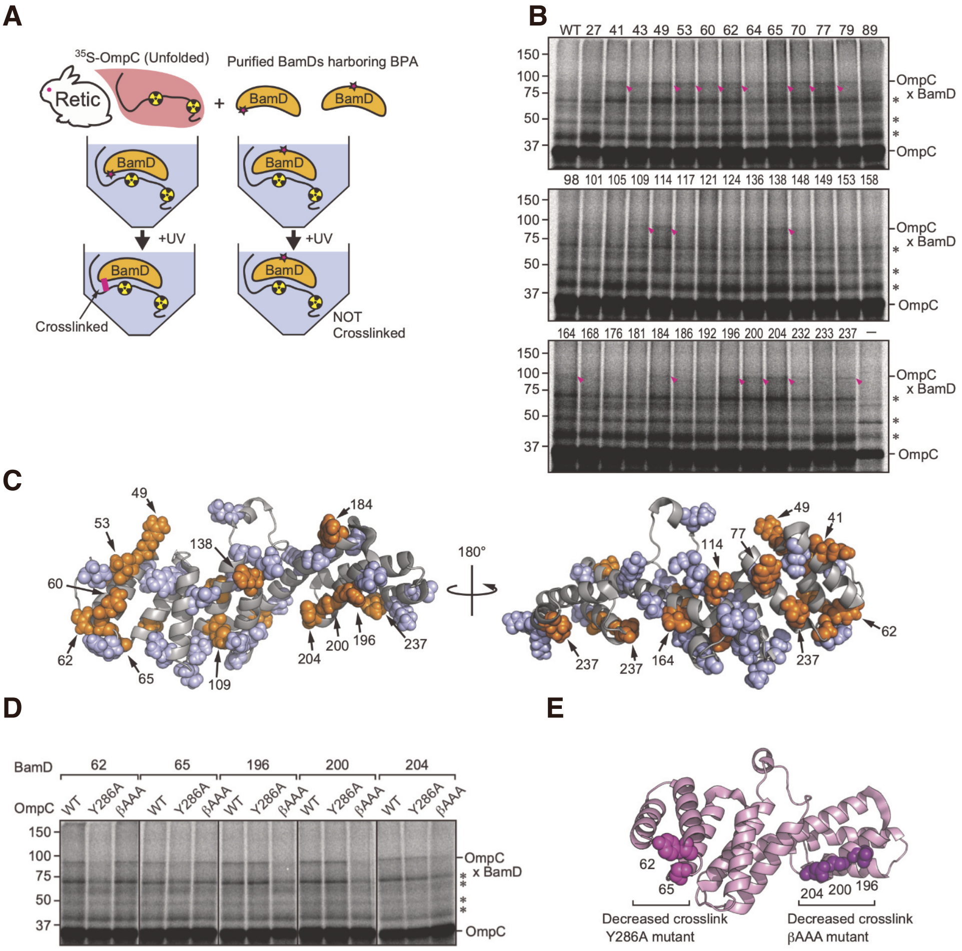
A
35S-OmpC (Unfolded)
Purified BamDs harboring BPA
Retic
BamD
+UV
Crosslinked
NOT
Crosslinked
B
WT 27 41 43 49 53 60 62 64 65 70 77 79 89
98 101 105 109 114 117 121 124 136 138 148 149 153 158
164 168 176 181 184 186 192 196 200 204 232 233 237 –
150
100
75
50
37
OmpC
x BamD
OmpC
C
180°
49 53 60 62 65 109 138 184 204 200 196 237
237 237 164 114 77 49 41 62
D
BamD
62 65 196 200 204
OmpC
WT Y286A βAAA
E
62
65
204 200 196
Decreased crosslink
Y286A mutant
Decreased crosslink
βAAA mutant

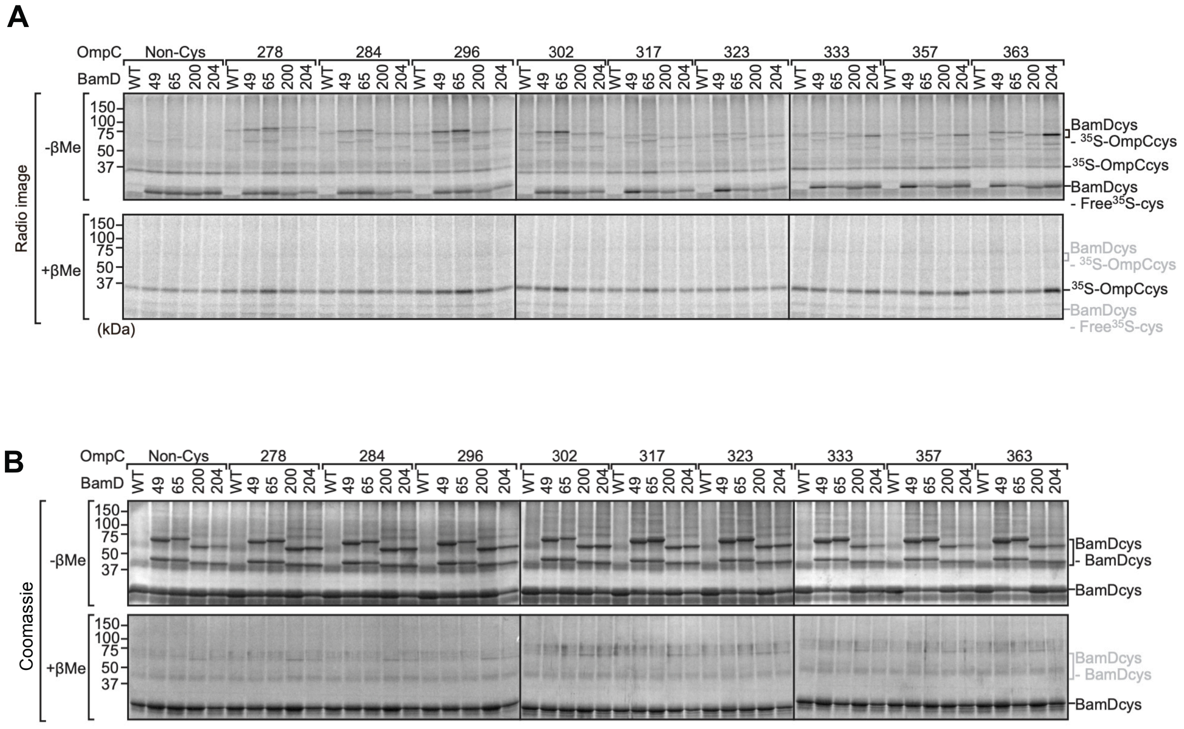
A
OmpC
Non-Cys
278
284
296
302
317
323
333
357
363
BamD
WT
49
65
200
204
Radio image
-βMe
+βMe
150
100
75
50
37
(kDa)
BamDcys
- 35S-OmpCcys
35S-OmpCcys
BamDcys
- Free35S-cys
B
Coomassie
BamDcys
- BamDcys
BamDcys

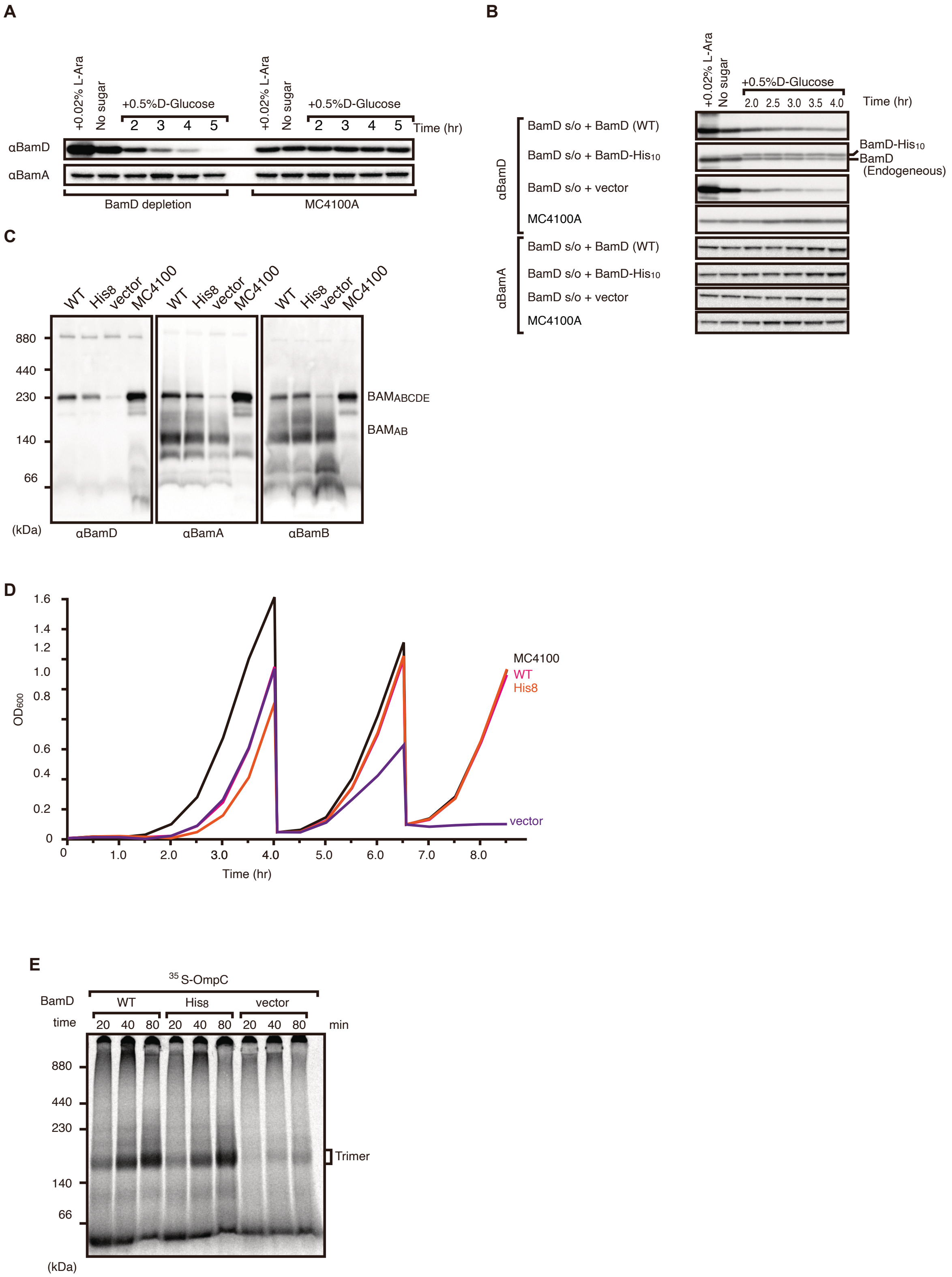
A
+0.02% L-Ara
No sugar
+0.5%D-Glucose
2 3 4 5
Time (hr)
αBamD
αBamA
BamD depletion
MC4100A
B
+0.02% L-Ara
No sugar
+0.5%D-Glucose
2.0 2.5 3.0 3.5 4.0
Time (hr)
BamD s/o + BamD (WT)
BamD s/o + BamD-His10
BamD s/o + vector
MC4100A
BamD-His10
BamD
(Endogeneous)
αBamD
αBamA
C
WT
His8
vector
MC4100
880
440
230
140
66
(kDa)
αBamD
αBamA
αBamB
BAMABCDE
BAMAB
D
OD600
1.6
1.4
1.2
1.0
0.8
0.6
0.4
0.2
0
1.0
2.0
3.0
4.0
5.0
6.0
7.0
8.0
Time (hr)
MC4100
WT
His8
vector
E
35 S-OmpC
BamD
WT
His8
vector
time
20 40 80 20 40 80 20 40 80
min
880
440
230
140
66
(kDa)
Trimer

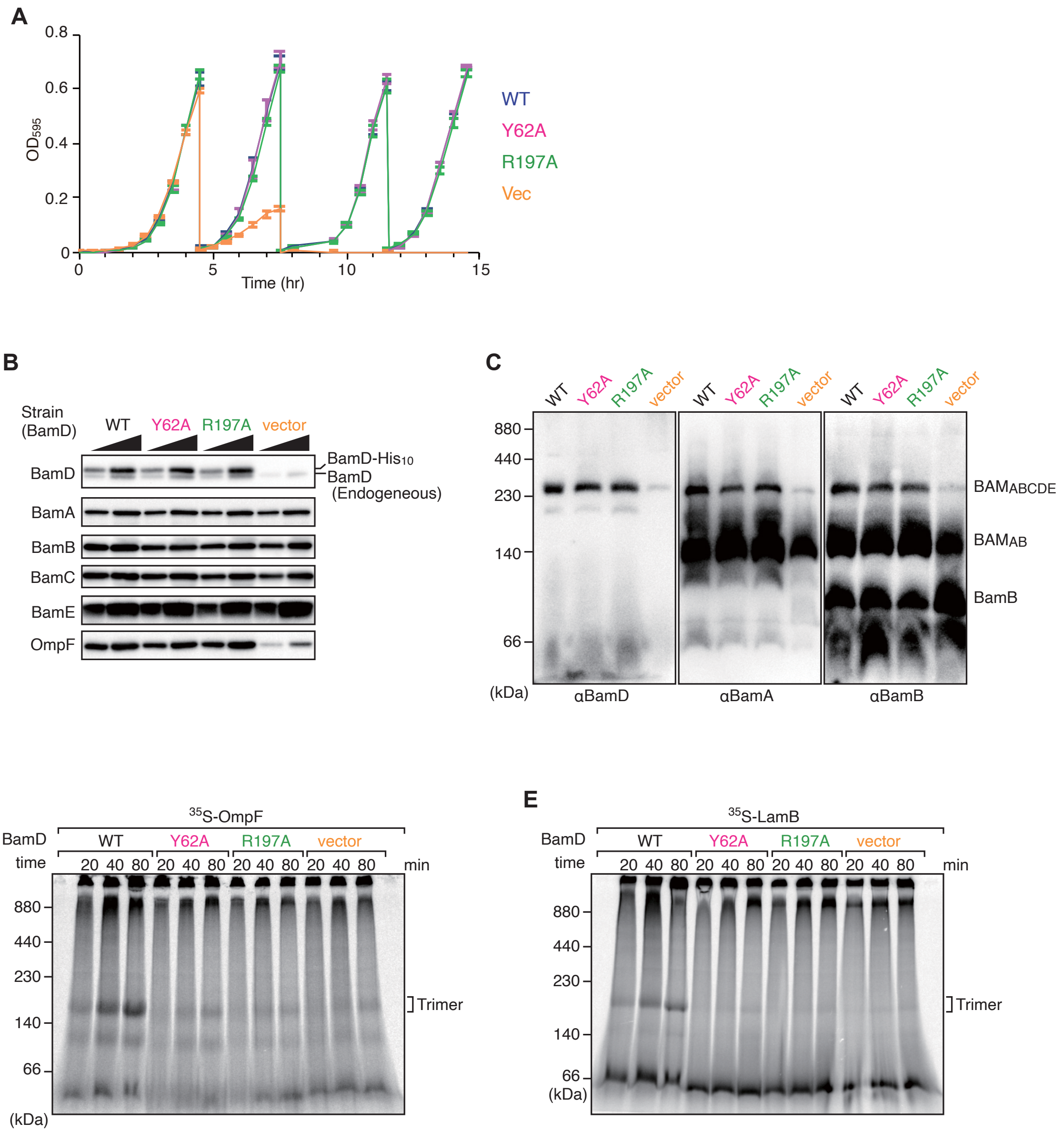
A
0.8
0.6
0.4
0.2
0
OD595
0
5
10
15
Time (hr)
WT
Y62A
R197A
Vec
B
Strain
(BamD)
WT
Y62A
R197A
vector
BamD
BamA
BamB
BamC
BamE
OmpF
BamD-His10
BamD
(Endogeneous)
C
WT
Y62A
R197A
vector
880
440
230
140
66
(kDa)
αBamD
αBamA
αBamB
BAMABCDE
BAMAB
BamB
D
35S-OmpF
BamD
time
WT
Y62A
R197A
vector
20 40 80 20 40 80 20 40 80 20 40 80
min
880
440
230
140
66
(kDa)
Trimer
E
35S-LamB
BamD
time
WT
Y62A
R197A
vector
20 40 80 20 40 80 20 40 80 20 40 80
min
880
440
230
140
66
(kDa)
Trimer

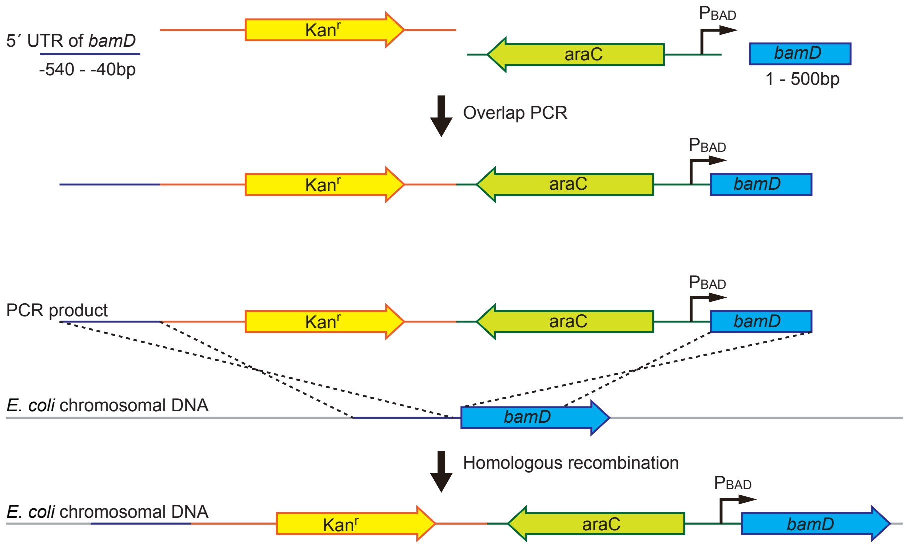

5´ UTR of bamD
-540 - -40bp
Kanr
araC
PBAD
bamD
1 - 500bp
Overlap PCR
Kanr
araC
PBAD
bamD
PCR product
Kanr
araC
PBAD
bamD
E. coli chromosomal DNA
bamD
Homologous recombination
PBAD
E. coli chromosomal DNA
Kanr
araC
bamD
