## Supplementary material for "Dual Recognition of Multiple Signals in Bacterial Outer Membrane Proteins Enhance Assembly and Maintain Membrane Integrity": Legends for Figure supplements

**Supplemental Figure 1. Schematic of *E. coli* microsomal membrane (EMM) isolation and EMM assembly assay.**

**(A)** *E. coli* microsomal membranes (EMM) were isolated via cell disruption by sonication. Cell debris is removed via low-speed centrifugation, finally membranes are harvested after high-speed centrifugation. **(B)** ^35^S-labelled EspP EMM assembly is performed as follows: (1) Radiolabeled substrate proteins are translated using rabbit reticulocytes lysate, (2) EMMs are treated with detergent then radio-labelled substrate is added to the EMM. (3) Substrate and BAM complex interacts with and assists the insertion of substrate into EMM. (4) EMMs were then incubated with protease. Unassembled EspP, as well as the passenger domain, are digested, only membrane-integrated barrel domain is protected from the protease. (5) Samples are boiled and subjected to SDS-PAGE for analysis.

**Supplemental Figure 2. Design of peptide inhibitor library from the *E* *coli* outer membrane protein OmpC.**

Primary sequence of the *E. coli* OMP OmpC divided into peptide regions. The sequence was divided into 23 peptides made up of 15 amino acids. Highlighted in the blue arrows are the 16 β-sheets which make up OmpC. Below OmpC sequence, the canonical β-signal of yeast-Tom40, a mitochondrial β-barrel protein, used as peptide #24 control.

**Supplemental Figure 3. Peptide inhibition of multiple BAM complex substrate assembly.**

**(A)** The 8 β-strand OMP, OmpA, was incubated with EMM then shifted onto ice to halt the assembly reaction at indicated times. Samples were split and incubated at either 25°C or 99°C for 10 min, before analysis by SDS-PAGE and radio-imaging. (u) and (f) indicate unfolded or folded, respectively. **(B-D)** Peptide inhibition of assembly reactions for OmpC (B), OmpF (C) and LamB (D). 35S-labelled proteins were incubated as above for indicated time, shifted onto ice, and solubilized in BN-PAGE lysis buffer containing 1.5% DDM. Solutions were clarified via centrifugation and analyzed by BN-PAGE and radio-imaging. The assembled, trimeric form of each porin is indicated. **(E)** Densitometry data for the endpoints of the assays on all assembled proteins (D-G), from 3 independent experiments.

**Supplemental Figure 4. Mutations to internal β-signal like motif results in loss of assembly in trimeric BAM complex substrate, OmpC.**

**(A)** Assembly of indicated BAM complex substrates with mutations to the -5 β-strand or final β-strand. Substrates were incubated with EMM for 10, 30, or 90 minutes, then analyzed by BN-PAGE and radio-imaging. Int. indicates high molecular weight intermediate form.

**Supplemental Figure 5. Multiple alignment of -5 β-strand of OMPs with 8, 10, or 12 β-strands.**

Sequence logos -5 and -3 β -strand were generated using WebLogo 3. Sequence from structure used is shown below the WebLogo. Aromatic, hydrophobic, positively charged, negatively charged, hydrophilic and secondary structure breaker residues (P, G) are shown by orange, green, blue, red, sky blue and black, respectively. The -5 and -3 strands are highlighted in pink with. PDB access codes of proteins are as follows: 8 β-strand; OmpA (1BXW), OmpW (2F1T), OmpX (1QJ8), PagP (3GP6), 10-strand; OmpT (1I78), 12 β-strand; EspP (2QOM), HBP (3AEH), NanC (2WJQ), PldaA/OmpLA (1QD5), Tsx (1TLW).

**Supplemental Figure 6. Multiple alignment of -5 β-strand of OMPs with 14, 16, or 18 β-strands.**

Sequence logos of -5 and -3 β-strand of were generated and colored as explained in Figure S3. PDB access codes of proteins are as follows: 14 β-strand; FadL (1T16) 16 β -strand; OmpC (2J1N), OmpF (1BT9), PhoE (1PHO), 18 β -strand; LamB (1MPM).

**Supplemental Figure 7. Multiple alignment of -5 β-strand of OMPs with 22 or 24 β-strands.**

Sequence logos of -5 and -3 β-strand of were generated and colored as explained in Figure S3. PDB access codes of proteins are as follows: 22 β-strand; BtuB (1NQE), CirA (2HDI), FhuA (1BY3), FepA (1FEP), FecA (1KMO), 24 β-strand; PapC (2VQI).

**Supplemental Figure 8. *In vitro* folding of OmpC β-signal mutants into detergent micelles.**

**(A)** Aromatic residues at the position C-terminus, not the 0Φ position, of the beta signal influences the folding rate of OmpC in vitro. Urea-denatured OmpC variants were folded into DDM-micelles in 10 mM phosphate buffer (pH 8.0), 2 mM EDTA, and 1 mM glycine. Samples were incubated at 37°C for up to 24 hours then split with half the samples **(B)** boiled and half at 25°C. OmpC showed three individual species, a high molecular weight assembled trimer, a middle molecular weight dimer and monomeric unfolded OmpC.

**Supplemental Figure 9. Ni-NTA pull-down of ^35^S-OmpC with purified BAM complex substrate binding subunits.**

**(A-C)** ^35^S-OmpC was incubated with purified BamB (A), BamC (), or BamD (bottom). Samples were rocked for 60 minutes at 4°C. Ni-NTA was added and samples were incubated at 4°C for an additional 30 minutes. Ni-NTA was pelleted by low speed centrifugation, flow through (FT) was collected and Ni-NTA was washed (W) with Buffer A containing 20 mM imidazole. BAM protein and substrate were eluted (E) with Buffer A and 400 mM imidazole. Proteins in FT, W, and E were precipitated using trichloroacetic acid, washed with acetone. Protein pellets were resuspended in SDS-loading dye, boiled, and resolved using SDS-PAGE. **(B)** ^35^S-OmpC Ni-NTA was incubated with BAM complex lipoproteins associated with substrate binding, BamB (top) and BamD (bottom). Experiment was performed as in (**A**).

**Supplemental Figure 10. Characterization of BamA-POPC, BamA-POPC-BamD, and BamA-POPC-BamD-OmpC (Y286A) interaction via NR.**

**(A)** Flow-chart of NR experiment. Chromium (Cr) and gold coated silicon-wafers were initially treated with Ni-NTA followed by the addition of purified His6-BamA. Lipid membrane, POPC, was add for the 1st measurement, after which BamD was incubated for 30 minutes for the 2nd measurement. Finally, unfolded OmpCY286A was added to the wafer for the 3rd measurement. **(B, D, F)** The neutron reflectometry profiles (symbols) and fits (lines) for BamA in the POPC bilayer **(B)**, the addition of BamD in the BamA-POPC **(D)**, and the addition of OmpC in the BamA-POPC-BamD complex **(F)** under D2O (black line), GMW (blue line) and H2O (red line) contrasts. **(C, E, G)** The real space nSLD profiles for BamA in the POPC bilayer **(C)**, the addition of BamD in the BamA-POPC **(E)**, and the addition of OmpC in the BamAPOPC-BamD complex **(G)** under D2O (black line), GMW (blue line) and H2O (red line) contrasts. **(D-E)** The addition of BamD, will not change the conformation of BamA and the volume fraction of BamD is found to be around 8.8% in P3-5 layer. **(F-G)** The further addition of OmpC (Y286A), will not change the conformation of BamAD complex and OmpC was located in the P3-5 layer with volume fraction of is 8.0%.

**Supplemental Figure 11. Characterization of BamA-POPC and BamA-POPC-BamD complex formation via NR.**

**(A)** NR profiles for BamA in membrane (1st measurement, black line), BamAD (2nd measurement, red line), and BamAD-OmpC (WT) (3rd measurement, blue line). **(B, D)** The neutron reflectometry profiles (symbols) and fits (lines) for BamA in the POPC bilayer **(B)** and the addition of BamD in the BamA-POPC **(D)** under D2O (black line) and GMW (blue line) contrasts. **(C, E)** The real space nSLD profiles for BamA in the POPC bilayer **(C)** and the addition of BamD in the BamA-POPC **(E)** under D2O (black line), GMW (blue line) and H2O (red line) contrasts. **(D, E)** The addition of BamD, did not change the conformation of BamA and the volume fraction of BamD was found to be around 8.8% in P3-5 layer. **(F)** Table summary of thickness, volume fraction, and roughness determined during the NR analysis. 1st, and 2nd displayed in black and magenta, respectively. P3-5: POTRA3, POTRA 4, and POTRA5, P1-2: POTRA1 and POTRA2, Lipid: POPC, Solution: D2O and gold match water (GMW).

**Supplemental Figure 12. *In vitro* BPA cross-link analysis of substrate binding regions within BamD.**

**(A)** Schematic of *in vitro* cross-linking of purified BamD and ^35^S-labelled OmpC. **(B)** BamD expressing unnatural amino acids were purified, incubated with purified OmpC substrate, UV-irradiated, and analyzed by SDS-PAGE. Numbers above each lane indicate the amino acid position that was mutated to BPA. Pink arrows designate the crosslink products of BamD-OmpC, while asterisks, signify non-specific cross-link products. **(C)** Summery of cross-link study in **(B)** on the structure of BamD. Positions highlighted in orange spheres indicate amino acid positions of BPA incorporation that cross-linked with OmpC, while positions which did not cross-link are highlighted in violet spheres. **(D)** Comparison cross-linking products of OmpC Wt, Y286A, and β-AAA (final β-strand) signal mutant and BamD. OmpC-BamD cross-link products are highlighted with a dash and non-specific cross-link products are as above. Experiment as preformed as in **(B)**. **(E)** Highlighted sites of BamD which showed decreased cross-linking products in OmpC signal mutations compared to Wt. Amino acids that showed decreased cross-linking with OmpC Y286A are shown in magenta (BamD 62 and 65), decreased cross-linking with OmpC β-AAA are shown in dark purple (BamD 196, 200, 204).

**Supplemental Figure 13. Analysis of the interaction between BamD and OmpC by intermolecular disulfide cross-linking.**

**(A)** Purified BamD^cys^ mutant proteins were incubated with ^35^S-OmpC with cysteine mutations from the -5 to the final β-strand. After the addition CuSO_4_ samples were purified with Ni-NTA and analyzed by SDS-PAGE and radio-imaging. Samples of the bottom gels were prepared in the presence of the reducing agent (β-mercaptoethanol; βMe). **(B)** Coomassie stained gels from **(A)**.

**Supplemental Figure 14. Validation and characterization of BamD depletion strain.**

**(A)** Optimization of BamD depletion as compared to parent-strain MC4100A. BamD levels were monitored by immunoblotting after growth in media containing glucose to repress chromosomal BamD expression as shown in Figure 6E. **(B)** Expression of plasmid-born BamD and BamD harboring poly-histidine tag versus empty vector and MC4100A strain. Both tagged and non-tagged BamD were detected at similar levels. **(C)** BamA expression remained unchanged between all samples. BN-PAGE Western blot analysis of BAM complex formation in BamD depletion strain. Expression of non-tagged and His-tagged BamD did not affect BAM complex formation. BamD depleted cells were unable to from full BAM complex (BamABCDE), but did form a smaller BamAB complex. **(D)** Growth curve of BamD depletion strain expressing His-tagged and non-tagged BamD as followed by optical density at 600 nm. Cells were grown in non-permissive conditions (+ glucose) and were sub-cultured into fresh LB media containing glucose two times, after which BamD depletion strain was incapable of growing. **(E)** Assembly of 35S-labelled OmpC in BamD depletion strain expressing plasmid-borne tagged and non-tagged BamD via EMM assembly assay. OmpC was able to be assembled by both tagged and non-tagged BamD, while BamD depleted cells were unable to assemble into OmpC into EMMs.

**Supplemental Figure 15. Effect of BamD mutations on cell growth and BAM complex formation *in vivo* and *in vitro* assembly ability.**

**(A**) Growth curve of BamD-depletion strain expressing BamD and mutant strains from plasmids. Cells were grown in glucose containing media to repress endogenous BamD expression. WT: blue line, Y62A: magenta, R197A: green, and empty vector: orange. **(B)** Steady-state levels of indicated proteins in EMM isolated from optimized time cultured bamD shut-down strains. BAM complex proteins showed no difference in steady state levels, while the OMP, OmpF, showed a decrease in empty vector. **(C)** BN-PAGE Western blot analysis of BAM complex formation in BamD depletion expressing mutant BamD. Both BamD mutations were able to form full BAM complex. **(D-E)** EMM assembly assay of BamD variants using OmpF **(D)** or LamB **(E)** as substrates, respectively. Assembly of both substrates were impaired in EMM containing BamD mutations.

**Supplemental Figure 16. Schematic model of BamD-depletion strain construction.**

Indicated four DNA fragments were amplified respectively, and then combined by overlap PCR method. The PCR fragment was introduced into MC4100A strain harboring pKD46 plasmid. Final chromosomal DNA map is shown at the bottom.
