## Supplemental tables for "Dual Recognition of Multiple Signals in Bacterial Outer Membrane Proteins Enhance Assembly and Maintain Membrane Integrity"

**Supplemental Table 1: Bacterial strains.**

| **Strain** | **Use** | **Ref/Source** |
| --- | --- | --- |
| BL21(DE3) | EMM isolation and *in vitro* studies. |  |
| BL21(DE3)* | Protein expression. | Invitrogen |
| MC4100A | *In vivo* studies, EMM isolation, and *in vitro* studies. |  |
| bamD-depletion | *In vivo* studies, EMM isolation, and *in vitro* studies. | This study |
| DH5α | Cloning strain. |  |

**Supplemental Table 2: Primers**

| **Primer name** | **Sequence 5'->3'** |
| --- | --- |
| pTnTl-f | ACTTAATACGACTCACTATAGGCTA |
| pTnT-r | GGATCCAAAAAACCCCTCAAGACCC |
| pSp64-f | ACCTTATGTATCATACACAT |
| pSp64-r | ACAGCTATGACATGATTACG |
| pTNTEspP-f | tgttctttttgcactcgagATGGATGTAACGCCGGTCATTAC |
| pTNTEspP-r | ccgcccgggtcgactctagaTCAGAACGAGTAACGGAAAT |
| pET15bOmpC-f | tgccgcgcggcagccatatgGAAGTTTACAACAAAGACGG |
| pET15bOmpC-r | ttagcagccggatcctcgagTTAGAACTGGTAAACCAGAC |
| pTnTOmpA-f | tgttctttttgcactcgagATGGCTCCGAAAGATAACACCTGGTACACTG |
| pTnTOmpA-r | ccgcccgggtcgactctagaTTAGAACTGGTAAACGATACCCACAGCAAC |
| pTnTOmpF-f | tgttctttttgcactcgagATGATGAAGCGCAATATTCTGGCAGTGATCG |
| pTnTOmpF-r | ccgcccgggtcgactctagaTTAGAACTGGTAAACGATACCCACAGCAACGGTGT |
| pTnTLamB-f | tgttctttttgcactcgagATGGTTGATTTCCACGGCTATGCACGTTCCGGTATTGGTTG |
| pTnTLamB-r | ccgcccgggtcgactctagaTTACCACCAGATTTCCATCTGGGCA |
| OmpCW77A-f | GACCGGTTACGGCCAGgcaGAATATCAGATCCAGG |
| OmpCW77A-r | CCTGGATCTGATATTCtgcCTGGCCGTAACCGGTC |
| OmpCE78A-f | CGGTTACGGCCAGTGGgcaTATCAGATCCAGGGCA |
| OmpCE78A-r | TGCCCTGGATCTGATAtgcCCACTGGCCGTAACCG |
| OmpCE78K-f | CGGTTACGGCCAGTGGaaaTATCAGATCCAGGGCA |
| OmpCE78K-r | TGCCCTGGATCTGATAtttCCACTGGCCGTAACCG |
| OmpCY79A-f | TTACGGCCAGTGGGAAgcaCAGATCCAGGGCAACA |
| OmpCY79A-r | TGTTGCCCTGGATCTGtgcTTCCCACTGGCCGTAA |
| OmpCK276A-f | CCTGGGTTGGGCGAACgcaGCACAGAACTTCGAAG |
| OmpCK276A-r | CTTCGAAGTTCTGTGCtgcGTTCGCCCAACCCAGG |
| OmpCK276E-f | CCTGGGTTGGGCGAACgaaGCACAGAACTTCGAAG |
| OmpCK276E-r | CTTCGAAGTTCTGTGCttcGTTCGCCCAACCCAGG |
| OmpCF280A-f | GAACAAAGCACAGAACgcaGAAGCTGTTGCTCAGT |
| OmpCF280A-r | ACTGAGCAACAGCTTCtgcGTTCTGTGCTTTGTTC |
| OmpCF280V-f | AACAAAGCACAGAACgtgGAAGCTGTTGCTCAG |
| OmpCF280V-r | CTGAGCAACAGCTTCcacGTTCTGTGCTTTGTT |
| OmpCE281A-f | CAAAGCACAGAACTTCgcaGCTGTTGCTCAGTACC |
| OmpCE281A-r | GGTACTGAGCAACAGCtgcGAAGTTCTGTGCTTTG |
| OmpCE281K-f | CAAAGCACAGAACTTCaaaGCTGTTGCTCAGTACC |
| OmpCE281K-r | GGTACTGAGCAACAGCtttGAAGTTCTGTGCTTTG |
| OmpCY286A-f | CGAAGCTGTTGCTCAGgcaCAGTTCGACTTCGGTC |
| OmpCY286A-r | GACCGAAGTCGAACTGtgcCTGAGCAACAGCTTCG |
| OmpCP294A-f | CGACTTCGGTCTGCGTgcaTCCCTGGCTTACCTGC |
| OmpCP294A-r | GCAGGTAAGCCAGGGAtgcACGCAGACCGAAGTCG |
| OmpCY325A-f | GATGTTGGTGCTACCgcgTACTTCAACAAAAAC |
| OmpCY325A-r | GTTTTTGTTGAAGTAcgcGGTAGCACCAACATC |
| OmpCY365A-f | AGCTCTGGGTCTGGTTgccCAGTTCTAAGTCGA |
| OmpCY365A-r | TCGACTTAGAACTGggcAACCAGACCCAGAGCT |
| OmpCBsigAAA-f | CTGGGTCTGGTTgccgcggcgTAAGTCGACCCGGGC |
| OmpCBsigAAA-r | GCCCGGGTCGACTTAcgccgcggcAACCAGACCCAG |
| OmpFV279A-f | AACAAAACGCAAGACgcgCTGTTAGTTGCGCAA |
| OmpFV279A-r | TTGCGCAACTAACAGcgcGTCTTGCGTTTTGTT |
| OmpFY285A-f | CTGTTAGTTGCGCAAgcgCAGTTCGATTTCGGT |
| OmpFY285A-r | ACCGAAATCGAACTGcgcTTGCGCAACTAACAG |
| LamBV333A-f | ACCAAGTGGTGGACCgcgGGTATTCGCCCGATG |
| LamBV333A-r | CATCGGGCGAATACCcgcGGTCCACCACTTGGT |
| LamBY339A-f | GGTATTCGCCCGATGgcgAAGTGGACGCCAATC |
| LamBY339A-r | GATTGGCGTCCACTTcgcCATCGGGCGAATACC |
| FLAGOmpC1-f | aacaggaggaattaaccATGAAAGTTAAAGTACTGTC |
| FLAGOmpC1-r | cttatcatcatcatccttataatcAGCAGCGTTTGCTGCGCCTG |
| FLAGOmpC2-f | gattataaggatgatgatgataagGAAGTTTACAACAAAGACGG |
| FLAGOmpC2-r | TACCAGCTGCAGATCTCGAGtcgacTTAGAACTGGTAAACCAGAC |
| BamDR49C-f | CAGGACGGTAACTGGtgcCAGGCAATAACGCAA |
| BamDR49C-r | TTGCGTTATTGCCTGgcaCCAGTTACCGTCCTG |
| BamDN60C-f | CTGGAAGCGTTAGATtgcCGCTATCCGTTTGGT |
| BamDN60C-r | ACCAAACGGATAGCGgcaATCTAACGCTTCCAG |
| BamDG65C-f | AATCGCTATCCGTTTtgcCCGTATTCGCAGCAG |
| BamDG65C-r | CTGCTGCGAATACGGgcaAAACGGATAGCGATT |
| BamDG200C-f | GTTAACCGCGTAGAAtgcATGTTGCGCGACTAC |
| BamDG200C-r | GTAGTCGCGCAACATgcaTTCTACGCGGTTAAC |
| BamDR203C-f | GTAGAAGGCATGTTGtgcGACTACCCGGATACC |
| BamDR203C-r | GGTATCCGGGTAGTCgcaCAACATGCCTTCTAC |
| BamDD204C-f | GAAGGCATGTTGCGCtgcTACCCGGATACCCAG |
| BamDD204C-r | GAAGGCATGTTGCGCtgcTACCCGGATACCCAG |
| OmpCQ278C-f | TGGGCGAACAAAGCAtgcAACTTCGAAGCTGTT |
| OmpCQ278C-r | AACAGCTTCGAAGTTgcaTGCTTTGTTCGCCCA |
| OmpCA282C-f | GCACAGAACTTCGAAtgcGTTGCTCAGTACCAG |
| OmpCA282C-r | CTGGTACTGAGCAACgcaTTCGAAGTTCTGTGC |
| OmpCA284C-f | AACTTCGAAGCTGTTtgcCAGTACCAGTTCGAC |
| OmpCA284C-r | GTCGAACTGGTACTGgcaAACAGCTTCGAAGTT |
| OmpCP294C-f | GACTTCGGTCTGCGTtgcTCCCTGGCTTACCTG |
| OmpCP294C-r | CAGGTAAGCCAGGGAgcaACGCAGACCGAAGTC |
| OmpCL296C-f | GGTCTGCGTCCGTCCtgcGCTTACCTGCAGTCT |
| OmpCL296C-r | AGACTGCAGGTAAGCgcaGGACGGACGCAGACC |
| OmpCY298C-f | CGTCCGTCCCTGGCTtgcCTGCAGTCTAAAGGT |
| OmpCY298C-r | ACCTTTAGACTGCAGgcaAGCCAGGGACGGACG |
| OmpCQ300C-f | TCCCTGGCTTACCTGtgcTCTAAAGGTAAAAAC |
| OmpCQ300C-r | GTTTTTACCTTTAGAgcaCAGGTAAGCCAGGGA |
| OmpCK302C-f | GCTTACCTGCAGTCTtgcGGTAAAAACCTGGGT |
| OmpCK302C-r | ACCCAGGTTTTTACCgcaAGACTGCAGGTAAGC |
| OmpCK317C-f | GACGAAGATATCCTGtgcTATGTTGATGTTGGT |
| OmpCK317C-r | ACCAACATCAACATAgcaCAGGATATCTTCGTC |
| OmpCV319C-f | GATATCCTGAAATATtgcGATGTTGGTGCTACC |
| OmpCV319C-r | GGTAGCACCAACATCgcaATATTTCAGGATATC |
| OmpCA323C-f | TATGTTGATGTTGGTtgcACCTACTACTTCAAC |
| OmpCA323C-r | GTTGAAGTAGTAGGTgcaACCAACATCAACATA |
| OmpCT333C-f | AACAAAAACATGTCCtgcTACGTTGACTACAAA |
| OmpCT333C-r | TTTGTAGTCAACGTAgcaGGACATGTTTTTGTT |
| OmpCV335C-f | AACATGTCCACCTACtgcGACTACAAAATCAAC |
| OmpCV335C-r | GTTGATTTTGTAGTCgcaGTAGGTGGACATGTT |
| OmpCN357C-f | GGCATCAACACTGATtgcATCGTAGCTCTGGGT |
| OmpCN357C-r | ACCCAGAGCTACGATgcaATCAGTGTTGATGCC |
| OmpCL363C-f | ATCGTAGCTCTGGGTtgcGTTTACCAGTTCTAA |
| OmpCL363C-r | TTAGAACTGGTAAACgcaACCCAGAGCTACGAT |
| OmpCF367C-f | GGTCTGGTTTACCAGtgcTAAGTCGACCCGGGC |
| OmpCF367C-r | GCCCGGGTCGACTTAgcaCTGGTAAACCAGACC |

**Supplementary Table 3: Primers for BPA introduction into BamD**

| **X** | **BamD(X)ambFwd** | **BamD(X)ambRev** |
| --- | --- | --- |
| 27 | GGGGTCAAAGGAAGAAtagCCTGATAATCCGCCAA | TTGGCGGATTATCAGGctaTTCTTCCTTTGACCCC |
| 41 | TTACGCGACTGCACAAtagAAGCTGCAGGACGGTA | TACCGTCCTGCAGCTTctaTTGTGCAGTCGCGTAA |
| 43 | GACTGCACAACAAAAGtagCAGGACGGTAACTGGA | TCCAGTTACCGTCCTGctaCTTTTGTTGTGCAGTC |
| 49 | GCAGGACGGTAACTGGtagCAGGCAATAACGCAAC | GTTGCGTTATTGCCTGctaCCAGTTACCGTCCTGC |
| 53 | CTGGAGACAGGCAATAtagCAACTGGAAGCGTTAG | CTAACGCTTCCAGTTGctaTATTGCCTGTCTCCAG |
| 60 | ACTGGAAGCGTTAGATtagCGCTATCCGTTTGGTC | GACCAAACGGATAGCGctaATCTAACGCTTCCAGT |
| 62 | AGCGTTAGATAATCGCtagCCGTTTGGTCCGTATT | AATACGGACCAAACGGctaGCGATTATCTAACGCT |
| 64 | AGATAATCGCTATCCGtagGGTCCGTATTCGCAGC | GCTGCGAATACGGACCctaCGGATAGCGATTATCT |
| 65 | TAATCGCTATCCGTTTtagCCGTATTCGCAGCAGG | CCTGCTGCGAATACGGctaAAACGGATAGCGATTA |
| 70 | TGGTCCGTATTCGCAGtagGTGCAGCTGGATCTCA | TGAGATCCAGCTGCACctaCTGCGAATACGGACCA |
| 77 | GCAGCTGGATCTCATCtagGCCTACTATAAAAACG | CGTTTTTATAGTAGGCctaGATGAGATCCAGCTGC |
| 79 | GGATCTCATCTACGCCtagTATAAAAACGCCGATT | AATCGGCGTTTTTATActaGGCGTAGATGAGATCC |
| 89 | CGATTTGCCGTTAGCAtagGCTGCCATCGATCGTT | AACGATCGATGGCAGCctaTGCTAACGGCAAATCG |
| 98 | CGATCGTTTTATTCGCtagAACCCGACCCATCCGA | TCGGATGGGTCGGGTTctaGCGAATAAAACGATCG |
| 101 | TATTCGCCTTAACCCGtagCATCCGAATATCGATT | AATCGATATTCGGATGctaCGGGTTAAGGCGAATA |
| 105 | CCCGACCCATCCGAATtagGATTATGTCATGTACA | TGTACATGACATAATCctaATTCGGATGGGTCGGG |
| 109 | GAATATCGATTATGTCtagTACATGCGTGGCCTGA | TCAGGCCACGCATGTActaGACATAATCGATATTC |
| 114 | CATGTACATGCGTGGCtagACCAATATGGCGCTGG | CCAGCGCCATATTGGTctaGCCACGCATGTACATG |
| 117 | GCGTGGCCTGACCAATtagGCGCTGGATGACAGTG | CACTGTCATCCAGCGCctaATTGGTCAGGCCACGC |
| 121 | CAATATGGCGCTGGATtagAGTGCGCTGCAAGGGT | ACCCTTGCAGCGCACTctaATCCAGCGCCATATTG |
| 124 | GCTGGATGACAGTGCGtagCAAGGGTTCTTTGGCG | CGCCAAAGAACCCTTGctaCGCACTGTCATCCAGC |
| 136 | CGATCGTAGCGATCGCtagCCTCAACATGCACGAG | CTCGTGCATGTTGAGGctaGCGATCGCTACGATCG |
| 138 | TAGCGATCGCGATCCTtagCATGCACGAGCTGCGT | ACGCAGCTCGTGCATGctaAGGATCGCGATCGCTA |
| 148 | TGCGTTTAGTGACTTTtagAAACTGGTGCGCGGCT | AGCCGCGCACCAGTTTctaAAAGTCACTAAACGCA |
| 149 | GTTTAGTGACTTTTCCtagCTGGTGCGCGGCTATC | GATAGCCGCGCACCAGctaGGAAAAGTCACTAAAC |
| 153 | TTCCAAACTGGTGCGCtagTATCCGAACAGTCAGT | ACTGACTGTTCGGATActaGCGCACCAGTTTGGAA |
| 158 | CGGCTATCCGAACAGTtagTACACCACCGATGCCA | TGGCATCGGTGGTGTActaACTGTTCGGATAGCCG |
| 164 | GTACACCACCGATGCCtagAAACGTCTGGTATTCC | GGAATACCAGACGTTTctaGGCATCGGTGGTGTAC |
| 168 | TGCCACCAAACGTCTGtagTTCCTGAAAGATCGTC | GACGATCTTTCAGGAActaCAGACGTTTGGTGGCA |
| 176 | GAAAGATCGTCTGGCGtagTATGAATACTCCGTGG | CCACGGAGTATTCATActaCGCCAGACGATCTTTC |
| 181 | GAAATATGAATACTCCtagGCCGAGTACTATACAG | CTGTATAGTACTCGGCctaGGAGTATTCATATTTC |
| 184 | ATACTCCGTGGCCGAGtagTATACAGAACGTGGCG | CGCCACGTTCTGTATActaCTCGGCCACGGAGTAT |
| 186 | CGTGGCCGAGTACTATtagGAACGTGGCGCATGGG | CCCATGCGCCACGTTCctaATAGTACTCGGCCACG |
| 192 | AGAACGTGGCGCATGGtagGCCGTCGTTAACCGCG | CGCGGTTAACGACGGCctaCCATGCGCCACGTTCT |
| 196 | ATGGGTTGCCGTCGTTtagCGCGTAGAAGGCATGT | ACATGCCTTCTACGCGctaAACGACGGCAACCCAT |
| 200 | CGTTAACCGCGTAGAAtagATGTTGCGCGACTACC | GGTAGTCGCGCAACATctaTTCTACGCGGTTAACG |
| 204 | AGAAGGCATGTTGCGCtagTACCCGGATACCCAGG | CCTGGGTATCCGGGTActaGCGCAACATGCCTTCT |
| 232 | GATGAATGCGCAAGCTtagAAAGTAGCGAAAATCA | TGATTTTCGCTACTTTctaAGCTTGCGCATTCATC |
| 233 | GAATGCGCAAGCTGAAtagGTAGCGAAAATCATCG | CGATGATTTTCGCTACctaTTCAGCTTGCGCATTC |
| 237 | TGAAAAAGTAGCGAAAtagATCGCCGCAAACAGCA | TGCTGTTTGCGGCGATctaTTTCGCTACTTTTTCA |

**Supplementary Table 4: Plasmids for *in vitro* translation.**

| **Plasmid name** | **Synthesized protein** | **Vector** | **Primers for construct** | **RE Site** | **Template DNA, source, or method** |
| --- | --- | --- | --- | --- | --- |
| pTnT-EspP | EspP | pTnT | pTNTEspP-f / pTNTEspP-r | XhoI/XbaI | gblock,  SLiCE |
| pTnT-OmpA | OmpA | pTnT | pTnTOmpA-f / pTnTOmpA-r | XhoI/XbaI | K-12 gene,  SLiCE |
| pTnT-OmpC | OmpC | pTnT | -- | XhoI/XbaI | (Gunasinghe et al., 2018) |
| pTNT-OmpF | OmpF | pTnT | pTnTOmpF-f / pTnTOmpF-r | XhoI/XbaI | K-12 gene, SLiCE |
| pTnT-LamB | LamB | pTnT | pTnTLamB-f / pTnTLamB-r | XhoI/XbaI | K-12 gene,  SLiCE |
| pTnT-OmpC-W77A | OmpC W77A | pTnT | OmpCW77A-f / OmpCW77A-r |  | Quick change mutagenesis |
| pTnT-OmpC-E78A | OmpC E78A | pTnT | OmpCE78A-f / OmpCE78A-r |  | Quick change mutagenesis |
| pTnT-OmpC-E78K | OmpC E78K | pTnT | OmpCE78K-f / OmpCE78K-r |  | Quick change mutagenesis |
| pTnT-OmpC-Y79A | OmpC Y79A | pTnT | OmpCY79A-f / OmpCY79A-r |  | Quick change mutagenesis |
| pTnT-OmpC-K276E | OmpC K276E | pTnT | OmpCK276E-f / OmpCK276E-r |  | Quick change mutagenesis |
| pTnT-OmpC-F276A | OmpC F276A | pTnT | OmpCK276A-f / OmpCK276A-r |  | Quick change mutagenesis |
| pTnT-OmpC-F280A | OmpC F280A | pTnT | OmpCF280A-f / OmpCF280A-r |  | Quick change mutagenesis |
| pTnT-OmpC-F280V | OmpC F280V | pTnT | OmpCF280V-f / OmpCF280V-r |  | Quick change mutagenesis |
| pTnT-OmpC-E281A | OmpC E281A | pTnT | OmpCE281A-f / OmpCE281A-r |  | Quick change mutagenesis |
| pTnT-OmpC-E281K | OmpC E281K | pTnT | OmpCE281K-f / OmpCE281K-r |  | Quick change mutagenesis |
| pTnT-OmpC-Y286A | OmpC Y286A | pTnT | OmpCY286A-f / OmpCY286A-r |  | Quick change mutagenesis |
| pTnT-OmpC-P294A | OmpC P294A | pTnT | OmpCP294A-f / OmpCP294A-r |  | Quick change mutagenesis |
| pTnT-OmpC-Y325A | OmpC Y325A | pTnT | OmpCY325A-f / OmpCY325A-r |  | Quick change mutagenesis |
| pTnT-OmpC-Y365A | OmpC Y365A | pTnT | OmpCY365A-f / OmpCY365A-r |  | Quick change mutagenesis |
| pTnT-OmpC-b-sigAAA | OmpC G352A, L354A, F359A | pTnT | OmpCBsigAAA-f / OmpCBsigAAA-r |  | Quick change mutagenesis |
| pTnT-OmpC-A284C,L296C | OmpC A284C, L296C | pTnT | OmpCA284C-f/r, OmpCL296C-f/r |  | Quick change mutagenesis |
| pTnT-OmpC-A323C, V335C | OmpC A323C, V335C | pTnT | OmpCA323C-f/r, OmpCV335C-f/r |  | Quick change mutagenesis |
| pTnT-OmpC-A323C, V335C | OmpC A323C, V335C | pTnT | OmpCA323C-f/r, OmpCV335C-f/r |  | Quick change mutagenesis |
| pTnt-OmpC-T333C, L363C | OmpC T333C, L363C | pTnT | OmpCT333C-f/r, OmpCL363C-f/r |  |  |
| pTnT-OmpF-V279A | OmpF V279A | pTnT | OmpFV279A-f / OmpFV279A-r |  | Quick change mutagenesis |
| pTnT-OmpF-Y285A | OmpF Y285A | pTnT | OmpFY285A-f / OmpFY285A-r |  | Quick change mutagenesis |
| pTnT-LamB-V333A | LamB V333A | pTnT | LamBV333A-f / LamBV333A-r |  | Quick change mutagenesis |
| pTnT-LamB-Y339A | LamB Y339A | pTnT | LamBY339A-f / LamBY339A-r |  | Quick change mutagenesis |

**Supplementary Table 5: Plasmids for recombinant protein expression.**

| **Plasmid name** | **Expressed protein** | **Vector/Promoter** | **Primers for construct** | **RE site** | **Template DNA, source, or method** |
| --- | --- | --- | --- | --- | --- |
| pET22-BamAL4H8 | BamA His8 at loop4 | pET22b/ T7 | -- |  | (Ding et al., 2020) |
| pHIS-BamB | His6-TEVsite-BamB | pHIS2-parallel/ T7 | -- |  | (Chen et al., 2021) |
| pHIS-BamC | His6-TEVsite-BamC | pHIS2-parallel/ T7 | -- |  | (Chen et al., 2021) |
| pHIS-BamD | His6-TEVsite-BamD | pHIS2-parallel/ T7 | -- |  | (Chen et al., 2021) |
| pET15b-OmpC | His6-TEVsite-OmpC | pET15b/ T7 | pTnTOmpC-f / pTnTOmpC-r | XhoI/XbaI | K-12 gene,  SLiCE |
| pET15b-OmpC Y286A | His6-TEVsite-BamD | pET15b/ T7 | OmpCY286A-f / OmpCY286A-r |  | Quick change mutagenesis |
| pBAD-FLOmpC | FLAG-OmpC-FLAG | pBAD/ areBAD | FLAGOmpC1-f /FLAGOmpC1-r, FLAGOmpC2-f /FLAGOmpC2-r |  | K-12 gene,  SLiCE |
| pBAD-FLOmpC F280A | FLAG-OmpC F280A | pBAD/ areBAD | OmpCF280A-f / OmpCF280A-r |  | pBAD-FLOmpC,  Quick change mutagenesis |
| pBAD-FLOmpC Y286A | FLAG-OmpC Y286A | pBAD/ areBAD | OmpCY286A-f / OmpCY286A-r |  | pBAD-FLOmpC,  Quick change mutagenesis |
| pBAD-FLOmpC F280A Y286A | FLAG-OmpC F280A Y286A | pBAD/ areBAD | OmpCY286A-f / OmpCY286A-r |  | pBAD-FLOmpC F280A  Quick change mutagenesis |
| pBAD-FLOmpC-VY | OmpC-FLAG V359A, Y365A | pBAD/ araBAD | OmpCV359A-f/r, OmpCY365A-f/r |  | pBAD-FLOmpC, quick change mutagenesis |
| pHIS-BamD (X) amber | His6-TEVsite-BamD (X) BPA | pBAD/ areBAD | BamD(X)ambFwd/ BamD(X)ambRev |  | pHIS-BamD,  Quick change mutagenesis |

**Supplementary Table 6: Plasmids for *in vivo* protein expression.**

| **Plasmid name** | **Expressed protein** | **Vector/Promoter** | **Primers for construct** | **RE site** | **Template DNA, source, or method** |
| --- | --- | --- | --- | --- | --- |
| pAp-BamD | BamD | pTnT/ BamA | BamDSL-f / BamDSL-r | XbaI/SalI | K-12 gene |
| pAp-BamD-His8 | BamD-His8 | pTnT/ BamA | BamDSL-f / BamDCHis8SL-r | XbaI/SalI | K-12 gene |
| pAp-BamD-His8 Y62A | BamD-His8 Y62A | pTnT/ BamA | BamDY62A-f / BamDY62A-r |  | pAp-BamD-His8, Quick change mutagenesis |
| pAp-BamD-His8 R197A | BamD-His8 R197A | pTnT/ BamA | BamDR197A-f / BamDR197A-r |  | pAp-BamD-His8, Quick change mutagenesis |

**Supplementary Table 7: Structural Data of this study**

| **Protein Name** | **PDB ID** | **PDB DOI** | **Citation** |
| --- | --- | --- | --- |
| OmpX | 1QJ8 | 10.2210/pdb1QJ8/pdb | (Vogt and Schulz, 1999) |
| OmpA | 1BXW | 10.2210/pdb1BXW/pdb | (Pautsch and Schulz, 1998) |
| OmpW | 2F1T | 10.2210/pdb2F1T/pdb | (Hong et al., 2006) |
| PagP | 3GP6 | 10.2210/pdb3GP6/pdb | (Cuesta-Seijo et al., 2010) |
| OmpT | 1I78 | 10.2210/pdb1I78/pdb | (Vandeputte-Rutten et al., 2001) |
| EspP | 2QOM | 10.2210/pdb2QOM/pdb | (Barnard et al., 2007) |
| HBP | 3AEH | 10.2210/pdb3AEH/pdb | (Tajima et al., 2010) |
| NanC | 2WJQ | 10.2210/pdb2WJQ/pdb | (Wirth et al., 2009) |
| PldA/OmpLA | 1QD5 | 10.2210/pdb1QD5/pdb | (Snijder et al., 1999) |
| Tsx | 1TLW | 10.2210/pdb1TLW/pdb | (Ye and van den Berg, 2004) |
| FadL | 1T16 | 10.2210/pdb1T16/pdb | (van den Berg et al., 2004) |
| OmpC | 2J1N | 10.2210/pdb2J1N/pdb | (Baslé et al., 2006) |
| PhoE | 1PHO | 10.2210/pdb1PHO/pdb | (Cowan et al., 1992) |
| OmpF | 1BT9 | 10.2210/pdb1BT9/pdb | (Phale et al., 1998) |
| LamB | 1MPM | 10.2210/pdb1MPM/pdb | (Dutzler et al., 1996) |
| BtuB | 1NQE | 10.2210/pdb1NQE/pdb | (Chimento et al., 2003) |
| CirA | 2HDI | 10.2210/pdb2HDI/pdb | (Buchanan et al., 2007) |
| FhuA | 1BY3 | 10.2210/pdb1BY3/pdb | (Locher et al., 1998) |
| FepA | 1FEP | 10.2210/pdb1FEP/pdb | (Buchanan et al., 1999) |
| FecA | 1KMO | 10.2210/pdb1KMO/pdb | (Ferguson et al., 2002) |
| PapC | 2VQI | 10.2210/pdb2VQI/pdb | (Remaut et al., 2008) |
